# ADCK4 deficiency destabilizes the coenzyme Q complex, which is rescued by 2,4-dihydroxybenzoic acid treatment

**DOI:** 10.1101/712323

**Authors:** Eugen Widmeier, Seyoung Yu, Anish Nag, Youn Wook Chung, Makiko Nakayama, Hannah Hugo, Florian Buerger, David Schapiro, Won-Il Choi, Jae-woo Kim, Ji-Hwan Ryu, Min Goo Lee, Catherine F. Clarke, Friedhelm Hildebrandt, Heon Yung Gee

## Abstract

*ADCK4* mutations usually manifest as steroid-resistant nephrotic syndrome, and cause coenzyme Q_10_ (CoQ_10_) deficiency. However, the function of ADCK4 remains obscure. We investigated ADCK4 function using mouse and cell models. Podocyte-specific *Adck4* deletion in mice significantly reduced survival and caused severe focal segmental glomerular sclerosis with extensive interstitial fibrosis and tubular atrophy, which were prevented by treatment with 2,4-dihydroxybenzoic acid (2,4-diHB), an analog of CoQ_10_ precursor molecule. ADCK4 knockout podocytes exhibited significantly decreased CoQ_10_ level, respiratory chain activity, mitochondrial potential, and dysmorphic mitochondria with loss of cristae formation, which were rescued by 2,4-diHB treatment, thus attributing these phenotypes to decreased CoQ_10_ levels. ADCK4 interacted with mitochondrial proteins including COQ5, and also cytoplasmic proteins including myosin and heat shock proteins. ADCK4 knockout decreased COQ complex levels, and the COQ5 level was rescued by ADCK4 overexpression in ADCK4 knockout podocytes. Overall, ADCK4 is required for CoQ_10_ biosynthesis and mitochondrial function in podocytes.

## Introduction

Coenzyme Q (CoQ, ubiquinone), a lipophilic component located in the inner mitochondrial membrane, Golgi apparatus, and cell membrane, plays a pivotal role in oxidative phosphorylation (Stefely & Pagliarini, 2017). CoQ shuttles electrons from complexes I and II to complex III in the mitochondrial respiratory chain (Mitchell, 1975). It also has a critical function in antioxidant defense owing to its redox potential (Mugoni, Postel et al., 2013). The CoQ biosynthesis pathway has been extensively studied in *Saccharomyces cerevisiae* (Tran & Clarke, 2007). At least 12 proteins encoded by the *Coq* genes form a complex, simultaneously stabilizing each other, and are involved in coenzyme synthesis (Tran & Clarke, 2007). Based on protein homology, approximately 15 homologous genes have been identified in humans (Stefely & Pagliarini, 2017).

Primary CoQ deficiencies due to mutations in ubiquinone biosynthetic genes (*COQ2, COQ4, COQ6, COQ7, COQ9, PDSS1, PDSS2, ADCK3*, and *ADCK4*) have been identified (Ashraf, Gee et al., 2013, Diomedi-Camassei, Di Giandomenico et al., 2007, Heeringa, Chernin et al., 2011, Lagier-Tourenne, Tazir et al., 2008, López, Schuelke et al., 2006, Mollet, Giurgea et al., 2007, Quinzii, Kattah et al., 2005). Clinical manifestations of CoQ_10_ deficiency vary depending on the genes involved, and mutations in the same gene can result in diverse phenotypes depending on the mutated allele (Acosta, Vazquez Fonseca et al., 2016). *COQ2 (Diomedi-Camassei et al., 2007), COQ6* (Heeringa et al., 2011), *PDSS2* (López et al., 2006), and *ADCK4* (Ashraf et al., 2013) have also been implicated in steroid-resistant nephrotic syndrome (SRNS). Although SRNS has no therapy, SRNS resulting from CoQ deficiency is unique because supplementation of CoQ_10_ alleviates the associated clinical symptoms (Ashraf et al., 2013, Heeringa et al., 2011, Korkmaz, Lipska-Zietkiewicz et al., 2016). This is partially true for ADCK4-related glomerulopathy, and several cases have been reported accordingly (Ashraf et al., 2013, Korkmaz et al., 2016). Recently, 2,4-dihydroxybenzoic acid (2,4-diHB), a metabolic intermediate of CoQ_10_, which can be used to bypass a defect in COQ7, has been shown to ameliorate disparate phenotypes in mouse models caused by heterogeneous enzymatic defects in CoQ biosynthesis (Wang, Oxer et al., 2015). Mutations in the *ADCK4* (aarF domain containing kinase 4, also known as *COQ8B*) gene generally manifest as adolescence-onset SRNS, sometimes accompanied with medullary nephrocalcinosis or extrarenal symptoms, including seizures (Ashraf et al., 2013, Park, Kang et al., 2017). The molecular mechanisms underlying SRNS resulting from *ADCK4* mutations are not well understood, largely because the function of ADCK4 is unclear.

ADCK3 (also known as COQ8A) and ADCK4 are two mammalian orthologs of yeast Coq8p/Abc1, which belongs to the microbial UbiB family; they appear to result from gene duplication in vertebrates (Kannan, Taylor et al., 2007, Lagier-Tourenne et al., 2008). UbiB and Coq8p are required for CoQ biosynthesis in prokaryotes and yeast, respectively, and are speculated to activate an unknown mono-oxygenase in the CoQ biosynthesis pathway (Kannan et al., 2007). Coq8p, ADCK3, and ADCK4 are present in the matrix of the inner mitochondrial membrane (Pagliarini, Calvo et al., 2008, Tauche, Krause-Buchholz et al., 2008, Vazquez Fonseca, Doimo et al., 2018). Coq8p is essential for the organization of high molecular mass Coq polypeptide complex and for phosphorylated forms of the Coq3, Coq5, and Coq7 polypeptides that are involved in methylation and hydroxylation steps in CoQ biosynthesis (He, Xie et al., 2014, Tauche et al., 2008). Similarly, it has been shown that ADCK3 interacts with CoQ biosynthesis enzymes in a protein complex (complex Q) (Floyd, Wilkerson et al., 2016). ADCK3 lacks protein kinase activity in the trans form and exhibits ATPase activity through its KxGQ motif (Stefely, Licitra et al., 2016). It is not clear whether ADCK4 functions in a manner similar to that of ADCK3. Therefore, in this study, we investigated the function of ADCK4 using mouse and cell models.

## Results

### Podocyte-specific *Adck4* knockout mice developed progressive proteinuria and severe focal segmental glomerular sclerosis, and presented increased mortality in adulthood

To evaluate the role of *Adck4* in kidney function, we generated a transgenic *Adck4* (*Adck4^tm1a^*) mouse line, using embryonic stem cells obtained from EUCOMM (**Appendix Fig S1A**). Efficient targeting of the *Adck4* gene was confirmed by genotyping (**Appendix Fig S1B and C**). Whole body loss of *Adck4* in *Adck4^tm1a^* mouse was found to be lethal, which is consistent with the report of the International Mouse Phenotyping Consortium (IPMC; www.mousephenotype.org). To circumvent *Adck4^tm1a^* embryonic lethality, we generated podocyte-specific *Adck4* knockout (KO) mice *Adck4^tm1d^* or *Nphs2.Cre^+^;Adck4^flox/flox^*(hereafter referred to as *Adck4^ΔPodocyte^*), by crossing *Nphs2-Cre+* mouse with *Adck4^flox/flox^* mouse in which two loxP sites surround exons 5 and 6 in the *Adck4* gene. While young *Adck4^ΔPodocyte^* mice appeared grossly normal, increased morbidity (hunched posture and seedy fur) (**Appendix Fig S1D**) and a significantly increased mortality (**Fig 1A**) were observed in older (> 9 months old) *Adck4^ΔPodocyte^* mice compared with those in littermate controls. Necropsy of 10-month-old *Adck4^ΔPodocyte^* mice revealed pale and significantly small kidneys compared with those in littermate controls (**Appendix Fig S1E and F**), indicating that podocyte-specific deletion of *Adck4* causes structural and functional kidney defects in *Adck4^ΔPodocyte^* mice. To examine the renal function of *Adck4^ΔPodocyte^* mice, we performed serial urine and plasma analyses for 18 consecutive months. *Adck4^ΔPodocyte^* mice displayed the first significant decrease in plasma albumin level at 5 months of age (**Appendix Fig S1G**) and the first significant increase in albumin/creatinine ratio (18.81 fold, *p* = 0.0005) remained significant throughout the study period compared with those in littermate controls (**Fig 1B**). The increase over time in albuminuria was the maximum, up to 31.2-fold, in *Adck4^ΔPodocyte^* mice compared with that in littermate controls (**Fig 1B**). The onset of kidney function decline in *Adck4^ΔPodocyte^* mice was associated with a significant increase in plasma creatinine and plasma BUN levels at 7 months of age, progressing to chronic kidney disease, followed by renal failure and consequently death (**Appendix Fig S1H-J and Fig 1A**). Histological analysis of kidneys from *Adck4^ΔPodocyte^* mice at 10 months of age demonstrated severe global and focal segmental glomerular sclerosis (FSGS) with extensive interstitial fibrosis and tubular atrophy (**Fig 1C and Appendix Fig S1K**). To characterize the glomerular phenotype of *Adck4^ΔPodocyte^* mice, we quantified the number of sclerotic glomeruli in the non-treated mutant mice at 10 months of age and found that non-treated *Adck4^ΔPodocyte^* mice had significantly increased number of sclerotic glomeruli (mean 96.03%) compared with that in non-treated littermate controls (**Appendix Fig S2C**). In addition, we compared the number of filtration slit units per micrometer of basement membrane in the glomeruli of *Adck4^ΔPodocyte^* and wild type mice. The filtration slit frequency was significantly reduced in *Adck4^ΔPodocyte^* mice compared with that in wild type mice (**Appendix Fig S2E**). To characterize the molecular abnormalities in the glomeruli of *Adck4^ΔPodocyte^* mice, we analyzed the expression pattern of the slit diaphragm proteins podocin (**Fig 1D**) and nephrin (**Appendix Fig S3A**), basement membrane marker nidogen, and primary process marker synaptopodin (**Appendix Fig S4A**) by confocal microscopy in the kidneys of 10-month-old mice. Staining of various podocyte markers was significantly reduced in the glomeruli of *Adck4^ΔPodocyte^* mice compared with that of the control glomeruli (**Appendix Fig S3C, S4B, and S5A and B**), while the basement membrane marker nidogen showed a higher expression in these mice than in normal mice (**Fig 1D and Appendix Fig S4A**), demonstrating that *Adck4* function is required for podocyte maintenance and homeostasis. As glomerular sclerosis is associated with increased expression of fibrotic markers, we analyzed the expression of the fibrotic markers, collagen IV (**Appendix Fig S3A**), and αSMA (**Appendix Fig S6A**) in the kidneys of *Adck4^ΔPodocyte^* mice by confocal microscopy. Indeed, the kidneys of *Adck4^ΔPodocyte^* mice presented significantly increased expression of collagen IV (**Appendix Fig S3E**) and αSMA (**Appendix Fig S6B**) in the glomeruli, characteristic of glomerular fibrosis. To study the structural changes in the glomeruli of *Adck4^ΔPodocyte^* mice at the ultrastructural level, we performed transmission electron microscopy (TEM) using the kidney of 10-month-old *Adck4^ΔPodocyte^* mice. Consistent with its localization and function in the mitochondria, the podocytes in the glomeruli of *Adck4^ΔPodocyte^* mice appeared to contain abnormal, functionally impaired mitochondria characterized by hyperproliferation and increased size (**Fig 1E**), presumably to compensate for their defective energy metabolism. The results revealed the abnormal structure of glomeruli, severe foot process effacement, and disturbed podocyte morphology in *Adck4^ΔPodocyte^* mice (**Fig 1E**). Overall, the glomerular phenotype of podocyte-specific *Adck4* KO mice recapitulates the pathology of FSGS in humans resulting from *ADCK4* mutations.

**Fig 1.**
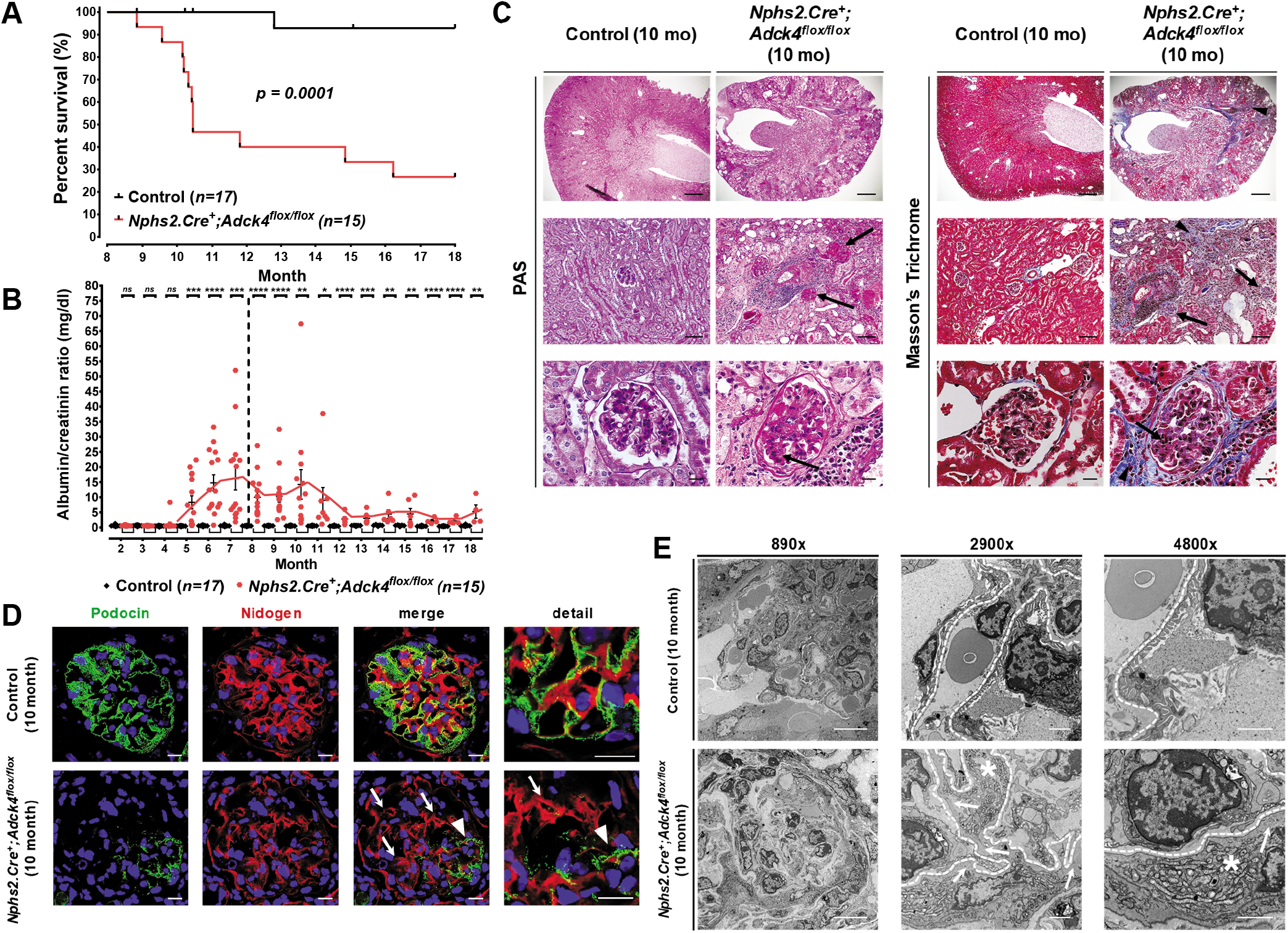
*Nphs2.Cre^+^;Adck4^flox/flox^* mice developed focal segmental glomerular sclerosis. (**A**) *Nphs2.Cre^+^;Adck4^flox/flox^* mutant mice exhibited reduced life span with a median survival period of 316 days and hazard ratio of 17.52 compared to that of littermate controls (Log-rank [Mantel–Cox] test, *p* = 0.0001; hazard Ratio [logrank]). (**B**) Urinary albumin/creatinine ratio serial analysis at indicated ages and genotypes revealed progressive proteinuria in *Nphs2.Cre^+^;Adck4^flox/flox^* mutant mice (red hexagon), but not in littermate controls (black diamond) (n = 15-17 animals per group). Dotted line displays the onset of renal failure. Note that once chronic renal failure ensues, urinary albumin excretion reduced as observed in SRNS (*p*-values were calculated using an unpaired *t*-test; *ns =* not significant, * *p* < 0.05, ** *p* < 0.01, *** *p* < 0.001, **** *p* < 0.0001; each data point represents the mean value of technical duplicates; the error bars represent SEM). (**C**) Kidney serial sections and representative images of 10-month-old mice. The *Nphs2.Cre^+^;Adck4^flox/flox^* mutant mice exhibited severe focal segmental glomerular sclerosis (arrows) with severe interstitial fibrosis and tubular atrophy (arrow heads). In contrast, wild type littermate control mice displayed normal histological kidney morphology (Scale bars: upper row 500 μm, middle row 100 μm, and lower row 20 μm). (**D**) Immunofluorescence staining for the slit diaphragm protein podocin (green) and the basement membrane marker nidogen (red). A normal expression pattern of podocin was observed in 10-month-old wild type littermate control mice. *Nphs2.Cre^+^;Adck4^flox/flox^* mutant mice mostly showed reduced podocin staining (arrows), appearing only on a few capillary loops (arrow head). (E) Transmission electron microscopy representative images of mice at the age of 10 months. *Nphs2.Cre^+^;Adck4^flox/flox^* mutant mice revealed severe podocyte foot process effacement (arrows) and an increased amount of dysmorphic mitochondria (asterisks). Glomerular basement membrane is highlighted by a dotted line (Scale bars: 10 μm left panel, and 2 μm middle and right panels).

### Treatment with 2,4-dihydroxybenzoic acid prevented the development of renal pathology in podocyte-specific *Adck4* knockout mice

Given that albuminuria started at around 4 months of age and renal structural abnormalities and functional decline manifested relatively late in *Adck4^ΔPodocyte^* mice, we decided to initiate the treatment with 2,4-diHB at 25 mM concentration in drinking water when the mice were 3 months old, in order to prevent the disease onset and to mitigate disease progression. Supplementation of 2,4-diHB did not have any effect on the survival rate, albumin/creatinine ratio, and kidney function and histology of the control mice (**Fig 2 A and B, and Appendix Fig S2, A and G-J**). *Adck4^ΔPodocyte^* mice treated with 2,4-diHB showed a normal survival rate despite maintaining proteinuria (**Fig 2B**) compared with that of healthy treated littermate controls (**Fig 2A**) and a significantly improved survival rate (p = 0.0078) compared with that of non-treated *Adck4^ΔPodocyte^* mice, which displayed an increased mortality rate progressing to end-stage renal disease (ESRD) with a median survival period of 316 days and hazard ratio of 9.36 (**Appendix Fig S2A**). The mortality rate reduction in *Adck4^ΔPodocyte^* mice treated with 2,4-diHB was associated with significantly improved plasma albumin level and renal function (**Appendix Fig S2, G to J**). This revealed normal glomerular histology (**Fig 2C and Appendix Fig S2B**) and a significantly reduced rate of sclerotic glomeruli (mean 14.76%) in 18-month-old treated *Adck4^ΔPodocyte^* mice (**Appendix Fig S2D**). The improvement in functional, histological, and ultrastructural findings in *Adck4^ΔPodocyte^* mice treated with 2,4-diHB was also associated with significantly improved expression of the slit diaphragm proteins podocin (**Fig 2D, and Appendix Fig S5C and D**) and nephrin (**Appendix Fig S3B and D**), although the expression of the primary process marker synaptopodin remained reduced (**Appendix Fig S4C and D**). Moreover, 18-month-old *Adck4^ΔPodocyte^* mice treated with 2,4-diHB showed significantly reduced expression of the fibrotic markers, collagen IV (**Appendix Fig S3B and F**) and αSMA (**Appendix Fig S6C and D**). Treatment with 2,4-diHB helped maintain the normal podocyte morphology and configuration at the ultrastructural level in the glomeruli of *Adck4^ΔPodocyte^* mice (**Fig 2E**) by preserving normal slit morphology, but the filtration slit frequency was decreased compared to that of littermate controls (**Appendix Fig S2F**). In summary, the treatment of *Adck4^ΔPodocyte^* mice with 2,4-diHB significantly prevented the development of FSGS and foot process effacement, maintaining normal renal function in treated mice at 18 months of age. Therefore, our findings demonstrate that 2,4-diHB is effective in protecting against renal disease progression and in improving survival in *Adck4^ΔPodocyte^* mice.

**Fig 2.**
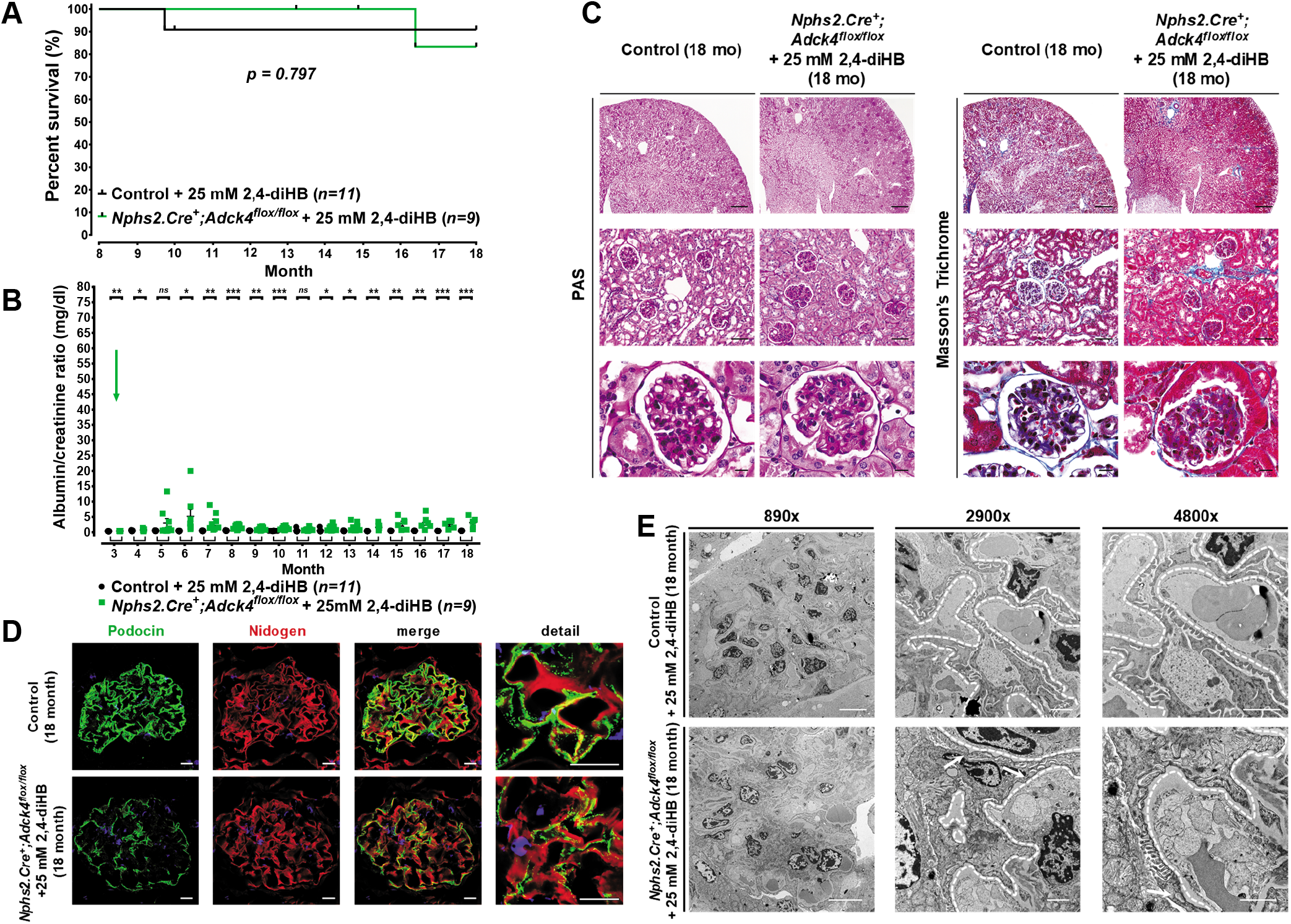
Treatment of *Nphs2.Cre^+^;Adck4^flox/flox^* mutant mice with 2,4-diHB prevented FSGS progression, resulting in normal survival rate. (**A**) *Nphs2.Cre^+^;Adck4^flox/flox^* mutant mice treated with 2,4-diHB presented similar survival rate as that of healthy treated littermate controls (Log-rank [Mantel–Cox] test, *p* = 0.797). (**B**) Urinary albumin/creatinine ratio serial analysis at indicated ages and genotypes (*n* = 9-11 animals per group). *Nphs2.Cre^+^;Adck4^flox/flox^* mutant mice treated with 2,4-diHB (green square) were protected from developing severe, progressive proteinuria, although proteinuria was significantly increased, compared with that in healthy treated littermate controls (black circle). Green arrow indicates the start of treatment (*p*-values were calculated using an unpaired *t*-test; *ns* = not significant, * *p* < 0.05, ** *p* < 0.01, *** *p* < 0.001; each data point represents the mean value of technical duplicates; the error bars represent SEM). (**C**) Kidney serial sections and representative images of 18-month-old mice. The wild type littermate control mice and *Nphs2.Cre^+^;Adck4^flox/flox^* mutant mice treated with 2,4-diHB displayed normal histological kidney morphology (Scale bars: upper row 500 μm, middle row 100 μm, and lower row 20 μm). (**D**) Immunofluorescence staining for the slit diaphragm protein podocin (green) and the basement membrane marker nidogen (red). The representative images of 18-month-old mice. A predominant expression of podocin was observed in *Nphs2.Cre^+^;Adck4^flox/flox^* mutant mice treated with 2,4-diHB compared with that in healthy treated wild type littermate control mice (Scale bars: 10 μm). (**E**) Transmission electron microscopy representative images of mice at the age of 18 months. In contrast, *Nphs2.Cre^+^;Adck4^flox/flox^* mutant mice treated with 2,4-diHB displayed mild foot process morphology changes with infrequent regions of effacement (arrows). Mitochondrial morphology remained normal (Scale bars: 10 μm left panel, and 2 μm middle and right panels)

### Loss of ADCK4 caused mitochondrial defects in podocytes

To investigate the function of ADCK4 at the cellular level, we generated *ADCK4* knockout (KO) cells of human podocytes and HK2 cells, which were originated from human proximal tubule epithelial cells. Each cell line was subjected to deletion of exon 6 of the *ADCK4* gene using the CRISPR/Cas9 system, and the absence of ADCK4 expression was confirmed by immunoblotting (**Appendix Fig S7A-E**). Knockout of ADCK4 did not affect the viability of both cell lines (**Appendix Fig S7F and G**). We have previously shown that the level of CoQ_10_ decreased in fibroblasts and lymphoblasts derived from patients with *ADCK4* mutations (Ashraf et al., 2013), but not in podocytes. Therefore, in the present study, we verified this finding using the established KO cells and found that ADCK4 KO resulted in decreased CoQ_9_ in both cultured podocytes and HK-2 cells compared to that in control cells (**Fig 3A**). However, CoQ_10_ was reduced only in cultured podocytes, but not in HK-2 cells (**Fig 3B**). The basal CoQ_10_ level in podocytes was three-fold higher than that in HK-2 cells (**Fig 3B**). As CoQ shuttles electrons from complexes I and II to complex III in the mitochondrial respiratory chain (Mitchell, 1975), the activity of complex II-III is dependent on the CoQ_10_ level in the mitochondria. Therefore, we measured the activity of complexes II and II-III and found that the activity of both was significantly reduced in ADCK4 KO podocytes (**Fig 3C**) compared to that in control cells, but not in ADCK4 KO HK-2 cells (**Fig 3D**). Decreased complex II-III activity observed in ADCK4 podocytes was partially rescued by the addition of 2,4-diHB to culture media (**Fig 3E**). The reduced form of CoQ (QH2) plays a role as a potent lipid-soluble antioxidant, scavenging free radicals and preventing lipid peroxidative damage (Stefely & Pagliarini, 2017). Although ADCK4 KO in itself did not affect the viability of cultured podocytes (**Appendix Fig S7F and G**), we examined cell viability upon arachidonic acid (AA) treatment because CoQ-deficient yeast mutants were found to be more sensitive to polyunsaturated fatty acids such as AA, which are prone to autoxidation and breakdown into toxic products (Do, Schultz et al., 1996). AA treatment reduced cell viability in both the control and ADCK4 KO podocytes, and ADCK4 KO podocytes were relatively more affected (**Fig 3F**). Decreased cell viability by AA treatment was rescued by supplementation of 2,4-diHB (**Fig 3F**). Overall, these findings suggested that the loss of ADCK4 caused CoQ deficiency and that podocytes were more susceptible than HK-2 cells were.

**Fig 3.**
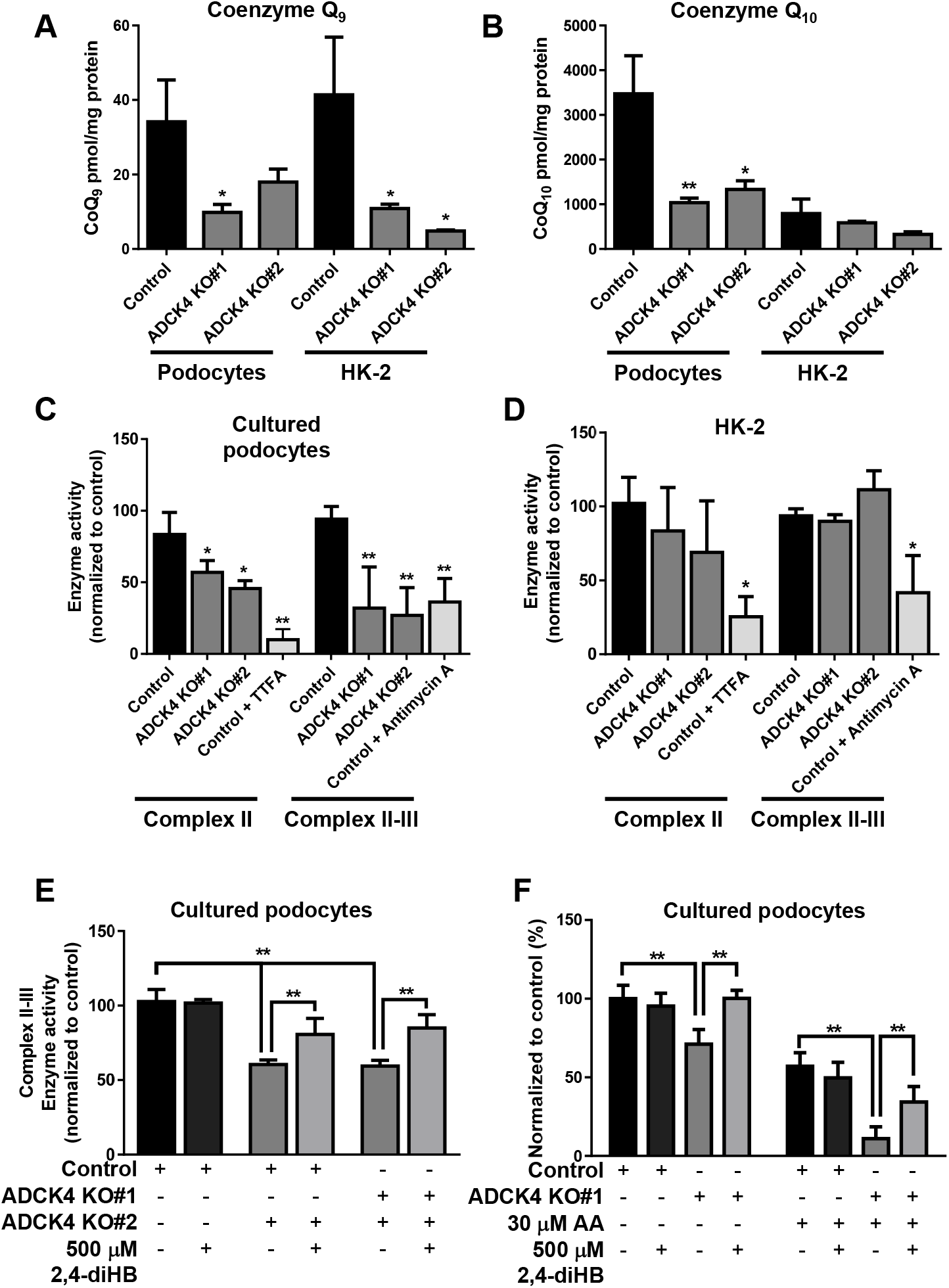
Coenzyme Q was deficient in ADCK4 knockout podocytes. (**A, B**) Coenzyme Q contents of cultured podocytes and HK-2 cells. The CoQ_9_ level was decreased in both cultured podocytes and HK-2 cells (A) and the CoQ_10_ level was severely deficient in ADCK4 KO podocytes (B). (**C, D**) Respiratory chain complex II and succinate-cytochrome c reductase (complex II-III) enzyme activities were measured in podocytes (C) and HK-2 cells (D). Complex II-III activities were decreased only in podocytes, whereas complex II activities were affected in both cell lines. (**E**) Succinate-cytochrome c reductase (complex II-III) enzyme activities were measured in podocytes. Decreased activities in ADCK4 KO podocytes were partially restored by the addition of 500 μM 2,4-dihydroxybenzoic acid (2,4-diHB). (**F**) Cell viability was measured using the Cell Count Kit-8 assay. ADCK4 KO podocytes exhibited susceptibility to 30 μM AA treatment. Decreased cell viability was reversed by the addition of 500 μM 2,4-diHB (*p*-values were calculated using an unpaired *t*-test; *ns* = not significant, * *p* < 0.05, ** *p* < 0.005; error bars represent mean ± SD).

To comprehensively understand the molecular changes induced by the KO of ADCK4, we performed proteomic analysis and quantified protein abundance changes by MS-based proteomics using isobaric tag for relative and absolute quantification (iTRAQ) (Chung, Lagranha et al., 2015) in podocytes with and without AA treatment. Proteomic characterization of the control and ADCK4 KO podocytes revealed more than 2,500 proteins, and 421 (16%) proteins were mitochondrial proteins. By Gene Ontology (GO) analysis, using DAVID functional annotation tool (david.abcc.ncifcrf.gov), differentially expressed proteins in the control and ADCK4 KO cells were divided into the following three categories of GO annotations: biological process, cellular component, and molecular function. The results indicated that proteins related to cellular defense response were upregulated and those associated with cytokine production pathway were downregulated in ADCK4 KO podocytes compared with those in the control cells (**Appendix Fig S7H**). In an injury situation, with the use of AA treatment, coenzyme metabolism-related proteins and intermediate filament related proteins were downregulated, whereas DNA regulation proteins were up-regulated in ADCK4 KO podocytes (**Appendix Fig S7I**).

### Disrupted mitochondrial morphology and mitochondrial membrane potential were observed in ADCK4 knockout podocytes

Abnormal proliferation of polymorphous mitochondria in the cytoplasm of podocytes is one of the characteristic ultrastructural findings of CoQ_10_-related diseases (Diomedi-Camassei et al., 2007). Similarly, patients with *ADCK4* mutations showed mitochondrial abnormalities in podocytes and proximal tubules (Park et al., 2017). We examined the ultrastructure of mitochondria in control and ADCK4 KO cells by TEM. The formation of cristae was disrupted and the shape of mitochondria was disintegrated in ADCK4 KO podocytes (**Fig 4A-H**), whereas ADCK4 KO HK-2 cells showed normal features of the mitochondria (**Appendix Fig S8A-H**). We also observed the effect of AA treatment on the ultrastructure of mitochondria by TEM. While the mitochondria of control podocytes were less affected by AA (**Fig 4I**), AA-treated ADCK4 KO podocytes showed more severe mitochondrial defects such as swollen and shortened cristae, and fewer inner membranes (**Fig 4J**). These results indicate that ADCK4 KO confers susceptibility to cellular stress, such as autoxidation of polyunsaturated fatty acids. To examine functional defects of mitochondria in ADCK4 KO cells, we measured the mitochondrial membrane potential using the JC-10 dye, which is concentrated in the mitochondrial matrix based on membrane polarization (Li, Yu et al., 2013). The results revealed the presence of inactive mitochondria in ADCK4 KO podocytes (**Fig 4K**). This finding was confirmed by another mitochondrial membrane potential assay using the potentiometric probe tetramethylrhodamine methyl ester (TMRM)-based fluorimetric assay. TMRM fluorescence intensity was significantly reduced in ADCK4 KO cells; however, supplementation of 2,4-diHB partially restored the lowered mitochondrial membrane potential (**Fig 4L**). In contrast, mitochondrial membrane potential was not different between the control and KO in HK-2 cells (**Appendix Fig S8I and J**). Therefore, the loss of ADCK4 caused morphological and functional defects of mitochondria in cultured podocytes by disrupting CoQ_10_ biosynthesis.

**Fig 4.**
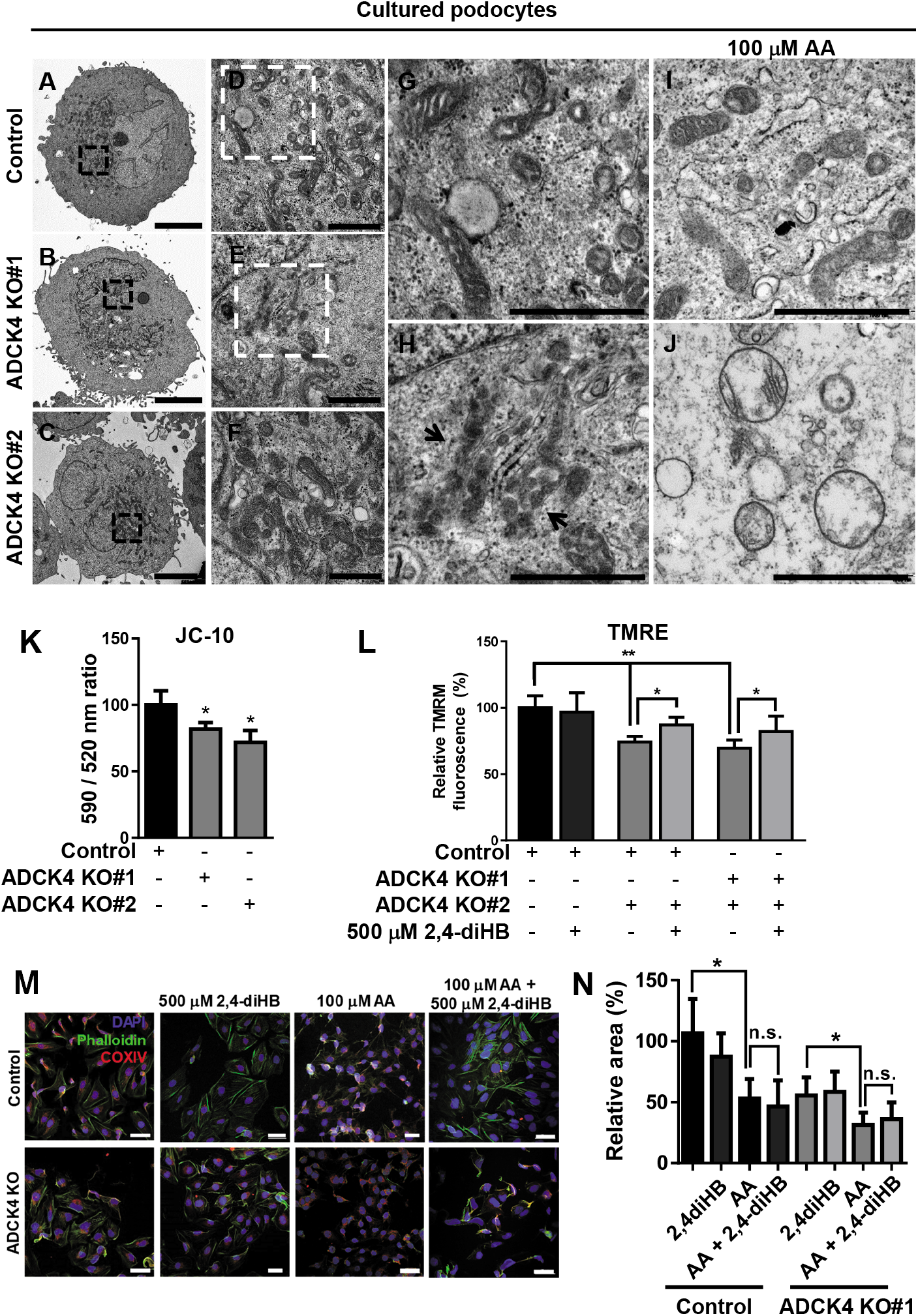
ADCK4 knockout podocytes showed mitochondrial defects. (**A-H**) TEM of podocytes showing mitochondrial morphology. Black (A and B) and white-boxed (D and E) areas are enlarged mitochondria in ADCK4 KO podocytes showing abnormal fission and disrupted cristae (H) (black arrows). (**I-J**) Mitochondria after AA treatment were severely disrupted in ADCK4 KO podocytes (J). (Scale bars: 0.5 μm in (A-C), 0.1 μm in (D-F), and 0.05 μm in (G-J)). (**K-L**) Mitochondrial membrane potential (ΔΨ) was measured using JC-10 (K) and TMRM (L). ADCK4 KO podocytes showed reduced ΔΨ compared with that of control podocytes. Reduced ΔΨ of ADCK4 KO podocytes was partially rescued by the addition of 500 μM 2,4-dihydroxybenzoic acid (2,4-diHB) (*p*-values were calculated using an unpaired *t*-test; *ns* = not significant, * *p* < 0.05, ** *p* < 0.005; the error bars represent mean ± SD). (**M**) Immunofluorescence of COXIV and phalloidin staining in podocytes. Phalloidin-stained area was shrunk in ADCK4 KO podocytes, and it became more prominent upon arachidonic acid (AA) treatment. Decreased cellular area was not reversed by the addition of 2,4-diHB. (Hundred cells from three independent experiments; *p*-values were calculated using an unpaired *t*-test; *ns* = not significant, * *p* < 0.05, ** *p* < 0.005; the error bars represent mean ± SD).

As we previously reported that ADCK4 knockdown reduced podocyte migration (Ashraf et al., 2013), in the present study, we examined the cytoskeleton of ADCK4 KO cells. Actin phalloidin staining revealed shrunk cellular area in ADCK4 KO podocytes (**Fig 4M**), whereas ADCK4 KO HK-2 showed surface area similar to that of control HK-2 cells (**Appendix Fig S8K and L**). Interestingly, shrunk cellular area in ADCK4 KO podocytes was not restored by 2,4-diHB supplementation, suggesting that this cellular phenotype is not related to decreased CoQ_10_ level and that ADCK4 might have other cellular functions in addition to its role in the CoQ biosynthesis pathway. AA treatment significantly reduced cellular area in both the control and ADCK4 KO podocytes (**Fig 4M**). Shrunk cellular area by AA treatment was not rescued by 2,4-diHB, further confirming that this phenotype is not related to CoQ_10_ deficiency (**Fig 4M and N**).

### ADCK4 interacted with and stabilized COQ proteins

We performed proteomic analysis to understand the function of ADCK4 via the identification of its interactome. We generated HEK293 cells that stably overexpressed C-terminal FLAG-tagged bacterial alkaline phosphatase (BAP; BAP-3xFLAG) and ADCK4 (ADCK4-3xFLAG) (**Appendix Fig S9A**). We confirmed that ADCK4-3xFLAG mostly localized to the mitochondria (**Appendix Fig S9B**). Following affinity purification using anti-FLAG beads (**Appendix Fig S9C and D**), protein eluates were analyzed using a liquid chromatograph coupled to a high-resolution mass spectrometer (LC-MS/MS). In total, 612 proteins were identified as interactors of ADCK4. Among them, the cytoplasmic proteins, including myosin (MYH10, MYH11, MYO1B, and MYO1C), filamin (FLNC), and kinase proteins (STK24, STK25, STK38 and ROCK1) were detected. In addition, the mitochondrial proteins, including ATP synthase subunit (ATP5L), cytochrome c oxidase subunit (COX6A1 and UQCRQ), and COQ5, were also identified as interactors (**Fig 5A**). GO analysis of mitochondrial interactors showed that these proteins are involved in transferase activity, oxidoreductase activity, nucleotide binding, and ATPase activity (**Fig 5B**). As COQ5, which functions as a C-methyltransferase in the CoQ biosynthesis pathway (Nguyen, Casarin et al., 2014), was identified as an interactor of ADCK4, we examined the COQ proteins in ADCK4 KO podocytes. Compared with that in the control podocytes, the expression of COQ3, COQ5, and COQ9 was significantly decreased in ADCK4 KO podocytes, indicating that these proteins in complex Q are destabilized in the absence of ADCK4 (**Fig 5C and D**). Decreased COQ5 was restored not only by transfection of wild type ADCK4, but also by 2,4-diHB (**Fig 5E**). In addition, we examined the effect of ADCK4 mutations, which were previously identified in individuals with nephrotic syndrome (Ashraf et al., 2013). COQ5 was also rescued by ADCK4 mutant proteins; however, the extent of the rescue was less than that by wild type ADCK4 (**Fig 5F**). In addition, as STK38 is a negative regulator of MAPKKK1/2 kinase signaling (Enomoto, Kido et al., 2008), we examined the MAPK pathway by western blotting and found that phosphorylated ERK1/2 was significantly increased in ADCK4 KO podocytes (**Fig 5G**). Moreover, we observed considerably enhanced MAPK signaling including p-p38, p-ERK1/2, and p-JNK under lipid peroxidation injury induced by AA (**Fig 5G**). As Coq2 silencing resulted in increased autophagy and mitophagy in *Drosophila* nephrocytes (Zhu, Fu et al., 2017), we examined mTOR and LC3 in ADCK4 KO podocytes; however, we could not observe increased LC3-II expression (**Appendix Fig S10**). Taken together, ADCK4 contributes to stabilizing the Q complex, elucidating CoQ deficiency in the absence of ADCK4, and other signaling pathways were also affected upon ADCK4 KO in podocytes.

**Fig 5.**
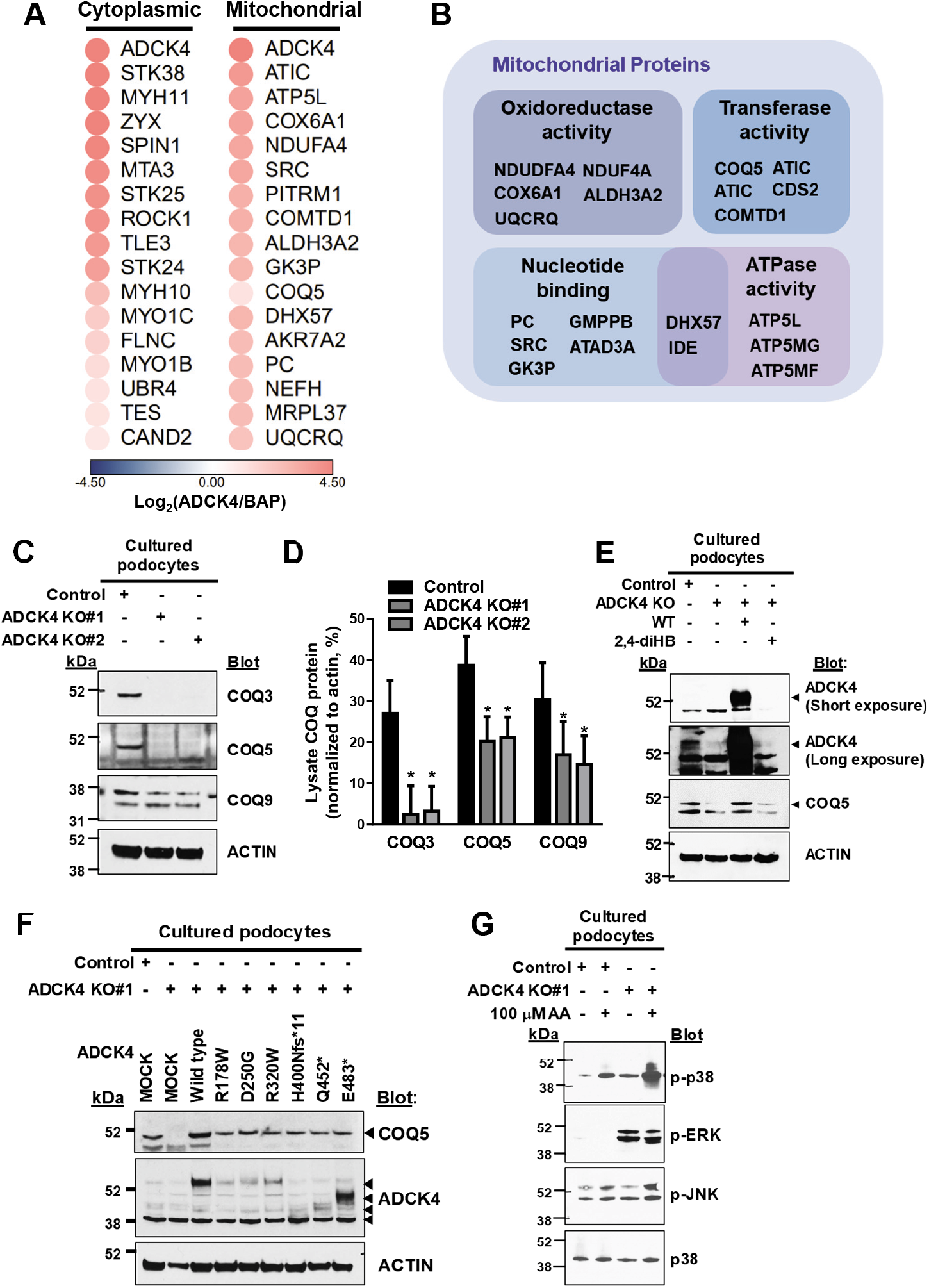
ADCK4 interacted with COQ5 and stabilized complex Q in podocytes. (**A**) ADCK4-interacting proteins isolated form HEK293 cells overexpressing ADCK4-3XFLAG. Both cytoplasmic and mitochondrial proteins were detected as interactors by NanoLC-MS/MS. (**B**) Gene ontology analysis showed that ADCK4-interacting mitochondrial proteins are associated with transferase activity, oxidoreductase activity, nucleotide binding, and ATPase activity. (**C**) Immunoblot showed that proteins in complex Q, COQ3, COQ5, and COQ9, were significantly reduced in ADCK4 knockout (KO) podocytes. (**D**) Densitometry analysis. Data are representative of at least three independent experiments and band intensities were normalized to that of β-actin. * *p* < 0.05; *t*-test. (**E**) Decreased COQ5 protein level was rescued by heterologous ADCK4 expression and 2,4-diHB treatment in ADCK4 KO podocytes. (**F**) Effect of ADCK4 mutations on COQ5 rescued in ADCK4 KO podocytes. COQ5 was rescued to a lower extent by truncated ADCK4 mutant proteins, whereas ADCK4 mutant proteins harboring missense variations were not different from wild type ADCK4. (**G**) Immunoblot analysis of the MAPK pathway. p-p38 and p-pERK were increased in ADCK4 KO podocytes and p-p38 was further induced by arachidonic acid (AA). Unphosphorylated p38 was used as the loading control.

## Discussion

In this study, we demonstrated that podocyte-specific deletion of *Adck4* in mice resulted in proteinuria and foot process effacement, recapitulating the features of nephrotic syndrome caused by *ADCK4* mutations. These defects were efficiently ameliorated by 2,4-diHB, an unnatural precursor analog of the CoQ biosynthesis pathway. ADCK4 KO podocytes exhibited reduced activity under complex II+III, mitochondrial defects, and sensitivity to AA, which resulted from CoQ deficiency. *ADCK4* mutations cause adolescence-onset nephrotic syndrome, which mostly progresses to ESRD in the second decade of life (Korkmaz et al., 2016). The late onset of renal disease is differentiated from nephropathy resulting from mutations in *WT1, NPHS1*, or *NPHS2*, which usually manifests in the first year of life. Similarly, in the present study, *Adck4^ΔPodocyte^* mice exhibited renal disease, which started around 4 months and progressed to ESRD by 12 months. ADCK4-associated glomerulopathy can be partially treated by CoQ_10_ supplementation, but it is not always successful (Ashraf et al., 2013, Feng, Wang et al., 2017, Korkmaz et al., 2016). Failure to respond to CoQ supplementation can be attributed to several possible causes, and one of the causes might be the progression of renal disease to the irreversible stage. Therefore, early genetic diagnosis is necessary to recognize *ADCK4* mutations. In this regard, as the onset of ADCK4-associated glomerulopathy is relatively late, if properly diagnosed, its therapy can be initiated before the disease becomes fulminant. Another reason for the failure of CoQ supplementation might be the poor oral availability of CoQ, which makes its therapeutic efficacy variable and limited. In the present study, we demonstrated that 2,4-diHB efficiently ameliorated proteinuria and prevented FSGS in *Adck4^ΔPodocyte^* mice. 2,4-diHB has also been shown to be more effective in tamoxifen-inducible *Mclk1/Coq7* KO mice than CoQ (Wang et al., 2015). It seems more readily absorbed than CoQ and is safe, as it has been used as a food flavor modifier owing to its sweet taste. Therefore, translational studies are required to investigate whether 2,4-diHB can be effective in individuals with *ADCK4* mutations.

2,4-diHB is expected to be beneficial for enzymatic deficiency in the CoQ biosynthetic pathway as it bypasses the defect in COQ7 (Stefely & Pagliarini, 2017, Wang et al., 2015). Therefore, it is actually interesting that 2,4-diHB rescued disease phenotypes in *Adck4^ΔPodocyte^* mice as ADCK4, although an uncharacterized mitochondrial protein, is not an enzyme directly involved in the CoQ biosynthetic pathway. This finding suggests that ADCK4 supports an enzymatic component in CoQ biosynthesis. We found that ADCK4 interacts with COQ5 by proteomic analysis and that the expression of proteins in complex Q, COQ3, COQ5, and COQ9, was significantly decreased in ADCK4 KO podocytes. It has been previously suggested that a physical and functional interaction between ADCK3 and COQ5 is important (Floyd et al., 2016, Stefely et al., 2016).

In the present study, phalloidin staining showed that the cytoskeleton of only podocytes was defective, but not that of HK-2 cells. This observation suggests that mitochondrial dynamics might also play important roles in maintaining the shape and function of podocytes because podocytes, like neurons (Imasawa & Rossignol, 2013), require a proper supply and high amount of energy to maintain their foot processes. In addition, as the foot process has rich microfilaments, these interactions might also participate in podocyte homeostasis (Greka & Mundel, 2012, Suleiman, Roth et al., 2017). Moreover, these cytoskeletal defects in ADCK4 KO podocytes were also consistent with the iTRAQ analysis data, which elucidated that cytoskeleton-related proteins were down-regulated in ADCK4 KO podocytes compared to those in control podocytes. In contrast to other cellular defects, the shrunk cytoskeleton of ADCK4 KO podocytes was not rescued by 2,4-diHB, suggesting that this cellular phenotype might not be related to CoQ_10_ deficiency. This also explains why CoQ

CoQ_10_ is well known for its antioxidant activity, protecting cells from oxidative stress (Mugoni et al., 2013). In this study, we treated the cells with AA, one of the polyunsaturated fatty acids, to induce lipid peroxidation stress and found that the MAPK pathway signaling was activated in ADCK4 KO podocytes. The MAPK signaling pathway is essential in regulating several cellular processes including inflammation, cell stress response, and cell proliferation (Cowan & Storey, 2003). In addition, AA treatment more significantly reduced cell viability in ADCK4 podocytes than in control podocytes, and the reduced viability was rescued by the supplementation of 2,4-diHB. The reduced form of CoQ_10_ might scavenge lipid peroxyl radicals and function as an antioxidant and prevent the initiation of lipid peroxidation as it has been reported to eliminate perferryl radicals (Ernster & Forsmark-Andree, 1993, Stefely & Pagliarini, 2017). In this regard, ADCK4 KO might confer hypersensitivity to lipid peroxidation stress. This suggests that cellular stress may be necessary to develop renal diseases in addition to loss of ADCK4, partially explaining the relative late onset of nephrotic syndrome resulting from *ADCK4* mutations.

Recent studies have revealed that ADCK3 lacks canonical protein kinase activity in the trans form; instead, it binds to lipid CoQ_10_ intermediates and exhibits ATPase activity(Reidenbach, Kemmerer et al., 2018, Stefely et al., 2016). In the present study, GO analysis also revealed that interactors of ADCK4, especially in the mitochondria, are significantly associated with oxidoreductase activity, which can be related to the antioxidant property of CoQ_10_. Furthermore, proteomic analysis revealed the ATP binding protein as an ADCK4 interactor, suggesting that ADCK4 has the ATPase activity, like ADCK3. Yet, the precise role of ADCK4 is not clear, and further studies are required to verify whether ADCK4 has ATPase or kinase activity towards an undiscovered substrate.

In conclusion, our study results suggest that ADCK4 in podocytes stabilizes proteins in complex Q in podocytes, and thereby contributes to CoQ synthesis and plays a role in maintaining the cytoskeleton structure. Cellular defects and renal phenotypes by ADCK4 deficiency were mostly rescued by 2,4-diHB supplementation, an unnatural precursor analog of CoQ_10_, demonstrating the important role of ADCK4 in the CoQ biosynthesis pathway. Our study provides insights into the functions of ADCK4 in CoQ biosynthesis and pathogenesis of nephrotic syndrome.

## Materials and Methods

### Mouse breeding and maintenance

The animal experimental protocols were reviewed and approved by the Institutional Animal Care and Use Committee of University of Michigan (#08619), Boston Children’s Hospital (#13-01-2283), and Yonsei University College of Medicine (#2015-0179). All mice were handled in accordance with the Guidelines for the Care and Use of Laboratory Animals. Mice were housed under pathogen-free conditions with a light period from 7:00 AM to 7:00 PM, and had *ad libitum* access to water and rodent chow. The mice were randomly assigned to the different experimental groups. Targeted *Adck4*^tm1a(EUCOMM)Hmgu^ (*Adck4*^tm1a^) embryonic stem cells were obtained from EUCOMM and injected into the blastocysts of mice. Chimeric mice were bred with C57BL/6J mice to establish germline transmission. *Nphs2.Cre^+^* (stock #008205) and *Pgk1.Flpo^+^* (#011065) mice were obtained from Jackson Laboratory. Genotyping was performed by standard PCR; the primer sequences are available upon request.

### Supplementation of 2,4-diHB to the mice in drinking water

2,4-diHB at a concentration of 25 mM was administered to the mice via drinking water and changed twice a week. The treatment was started at 3 months of age and continued up to 18 months of age.

### Urine analysis

Urine was collected by housing the mice overnight (12 h) in metabolic cages. All samples were immediately frozen and stored at −80°C. The samples were thawed on ice prior to urine albumin and creatinine measurements. Urinary albumin was measured using the Albumin Blue Fluorescent Assay Kit (Active Motif), as per the manufacturer’s instructions. Urine creatinine was measured using the LC-MS/MS method as described previously (Young, Struys et al., 2007) Proteinuria was expressed as milligram of albumin per milligram of creatinine.

### Whole blood and plasma analysis

Blood from the mice was drawn using the facial vein bleeding method and collected in citrate tubes. The whole blood sample was subsequently analyzed using the Vetscan^®^ VS2 Chemistry Analyzer, as per manufacturer’s instructions. Plasma samples obtained by centrifugation of whole blood were immediately frozen at −80°C. Plasma creatinine was measured using the LC-MS/MS method as described previously (Young et al., 2007).

### Immunoblotting and immunofluorescence staining

These experiments were performed as described previously (Hinkes, Wiggins et al., 2006). Anti-Podocin, anti-FLAG M2 (Sigma-Aldrich), anti-Nidogen (Novus), anti-Nephrin (Progen), anti-COQ3, anti-COQ5, anti-COQ9 (Proteintech), anti-p-mTOR, anti-mTOR, anti-p-p38, anti-p-ERK, anti-pJNK, anti-p38, anti-LC3 (Cell Signaling), anti-COXIV, anti-actin (Abcam), and ADCK4 (LSBio) were purchased from the indicated commercial sources. Alexa Fluor 488 Phalloidin and secondary antibodies were purchased from Invitrogen. Fluorescent images were obtained using an SP5X laser scanning microscope (Leica) or LSM 700 microscope (Carl Zeiss). Images were processed and analyzed using Leica AF, ImageJ, and Adobe Photoshop CS6 software.

### Histological analysis

The kidney tissues were fixed in 4% paraformaldehyde (PFA), sectioned (5 μm thickness), and stained with hematoxylin and eosin, periodic acid-Schiff, Masson’s trichrome, and Jone’s silver following the standard protocols for histological examination.

### Ultrastructural analysis

The kidney tissues and cells were fixed in 2.5% glutaraldehyde, 1.25% PFA, and 0.03% picric acid in 0.1 M sodium cacodylate buffer (pH 7.4) overnight at 4°C. They were washed with 0.1 M phosphate buffer, post-fixed with 1% OsO_4_ dissolved in 0.1M phosphate-buffered saline (PBS) for 2 h, dehydrated in ascending gradual series (50–100%) of ethanol, and infiltrated with propylene oxide. Samples were embedded using the Poly/Bed 812 Kit (Polysciences). After pure fresh resin embedding and polymerization in a 65°C oven (TD-700, DOSAKA, Japan) for 24 h, sections of approximately 200–250 nm thickness were cut and stained with toluidine blue for light microscopy. Sections of 70-nm thickness were double stained with 6% uranyl acetate (EMS, 22400) for 20 min and lead citrate (Fisher) for 10 min for contrast staining. The sections were cut using LEICA EM UC-7 (Leica) with a diamond knife (Diatome) and transferred on to copper and nickel grids. All the sections were observed by transmission electron microscopy (JEM-1011, JEOL) at an acceleration voltage of 80 kV.

### Plasmids, cell culture, transfection, and lentivirus transduction

sgRNAs targeting human ADCK4 (sgRNA1: GCTGCACAATCCGCTCGGCAT, sgRNA2: GTAAGGTCTGCACAATCCGCT, and sgRNA3: GACCTTATGTACAGTTCGAG,) were cloned into BsmBI-digested lentiCRISPR v2 (Addgene plasmid #52961). ADCK4 cDNA was cloned into the p3xFLAG CMV26 (C-terminal) vector (Sigma-Aldrich). BAP cDNA cloned into the p3xFLAG CMV7 vector was digested using Kpn1 and EcoR1 restriction enzymes (New England BioLabs) and ligated into the p3xFLAG CMV24 vector.

Immortalized human podocytes (Saleem, O’Hare et al., 2002) were maintained in RPMI + GlutaMAX™-I (Gibco) supplemented with 10% FBS, penicillin (50 IU/mL)/streptomycin (50 μg/mL), and insulin-transferrin-selenium-X. Human proximal tubule cells (HK-2) and HEK293 were maintained in DMEM supplemented with 10% FBS and 1% penicillin/streptomycin. Plasmids were transfected into podocytes or HEK293 cells using Lipofectamine 2000 (Invitrogen). HEK293 cells stably expressing p3xFLAG-ADCK4 or BAP were selected and maintained with 1 mg/mL G418.

To establish ADCK4 KO cells, lentiCRISPR v2, pMD2.G, and psPAX2 were transfected into Lenti-X 298T cells (Clontech). Supernatant containing lentivirus was collected 48 h after transfection and passed through a 0.2-M filter. Cultured podocytes and HK-2 cells were transduced with lentivirus, selected, and maintained with 4 μg/mL puromycin.

### Cell viability assay

Cell viability assay was performed using the Cell Counting Kit-8 (Dong-in bio.). Cell suspension (100 μL; 1 × 10^5^/mL) with culture medium was added to a 96-well plate and incubated for 24 h in a CO_2_ incubator. The medium was replaced with phenol-free fresh medium with or without 30 μM AA and/or 500 μM 2,4-diHB for 15 h. Four wells were included under the same conditions. CCK-8 reagent (10 μL) was added to each well and the cells were incubated for 1 h; optical density of the sample at 450 nm was measured.

### Cellular lipid extraction and CoQ measurements via HPLC-MS/MS

Cells (approximately 0.1 g) were thawed on ice and resuspended in 1.5 mL of PBS (0.14 M NaCl, 12.0 mM NaH_2_PO_4_, and 8.1 mM Na_2_HPO_4_; pH 7.4), followed by homogenization using a polytron (Kinematica PT 2500E) for 1 min at 10 000 rpm on ice. Lipid extracts were prepared as previously described with some minor modifications(Fernández-Del-Río, Nag et al., 2017). Briefly, dipropoxy-CoQ_10_ was used as the internal standard and was added at a constant volume to all the cell pellets and to a set of five CoQ_9_ and CoQ_10_ standards of known concentrations ranging linearly from 7.2 pmol to 400 pmol (to obtain a typical standard curve for CoQ quantification in the cell pellets). The samples were vortexed in 2 mL of methanol for 30 s, followed by addition of 2 mL petroleum ether. After vortexing for an additional 30 s, the organic upper layer was transferred to a new tube. Another 2 mL of petroleum ether was added to the original methanol layer, and samples were vortexed again for 30 s. The organic phase was removed, and the combined organic phase was dried under a stream of nitrogen gas. The samples were resuspended in 200 μL of ethanol containing 1 mg/mL benzoquinone to oxidize all the lipids. Chromatographic separation was achieved through a reverse phase Luna 5 μM PFP(2) column (Phenomenex) with a mobile phase comprised of 90% solvent A (95:5 mixture of methanol:isopropanol containing 2.5 mM ammonium formate) and 10% solvent B (isopropanol containing 2.5 mM ammonium formate) at a constant flow rate of 1 mL/min. Transitions monitored were: m/z 795.6/197.08 (CoQ_9_), m/z 812.6/197.08 (CoQ_9_ with ammonium adduct), m/z 863.6/197.08 (CoQ_10_), m/z 880.6/197.08 (CoQ_10_ with ammonium adduct), m/z 919.7/253.1 (dipropoxy-CoQ_10_), and m/z 936.7/253.1 (dipropoxy-CoQ_10_ with ammonium adduct).

### Mitochondrial respiratory enzyme activity measurement

Cell lysate (15–50 μg) was diluted in phosphate buffer (50 mM KH_2_PO_4_; pH 7.5), and then subjected to spectrophotometric analysis for isolated respiratory chain complex activities at 37°C using a spectrophotometer (PerkinElmer). Complex II activity was measured at 600 nm (ε = 19.1 mmol^-1^cm^-1^) after the addition of 20 mM succinate, 80 μM dichlorophenolindophenol (DCPIP), 300 μM KCN, and 50 μM decylubiquinone. Complex II activity was defined as the flux difference with or without 10 mM malonate. Complex II+III activity was also determined at 550 nm (ε = 18.5 mmol^-1^cm^-1^) in the presence of 10 mM succinate, 50 μM cytochrome c, and 300 μM KCN. Complex II-III activity was defined as the flux difference before and after the addition of 10 mM thenoyltrifluoroacetone (TTFA). All chemicals were obtained from Sigma-Aldrich.

### Isobaric tag labeling for relative and absolute quantification

Isobaric tag labeling for relative and absolute quantification (iTRAQ) was performed by Poochon Scientific as described previously (Chung et al., 2015). Proteins (100 μg) were extracted from control and ADCK4 KO podocytes, digested with trypsin, and labeled using the 8-plex iTRAQ Labeling Kit (AB Sciex). The fractionation of iTRAQ-multiplex labeled peptide mixture was carried out using Agilent AdvanceBio Column (2.7 μm, 2.1 × 250 mm) and Agilent UHPLC 1290 system (Agilent, Santa Clara, CA). The LC/MS/MS analysis was carried out using Thermo Scientific Q-Exactive hybrid Quadrupole-Orbitrap Mass Spectrometer and Thermo Dionex UltiMate 3000 RSLCnano System (Thermo, San Jose, CA). MS Raw data files were searched against the human protein sequence databases obtained from the NCBI website using Proteome Discoverer 1.4 software (Thermo) based on the SEQUEST and percolator algorithms.

### Identification of ADCK4 interactors

Proteins (75 mg) from HEK293 cells stably expressing p3XFLAG-ADCK4 or -BAP were incubated with 80 μL of FLAG M2 agarose beads (Sigma-Aldrich) for 48 h at 4°C in an orbital shaker. The agarose beads were washed four times with lysis buffer to restrict non-specific binding. Subsequently, 200 μL of elution buffer containing 150 ng/μL 3xFLAG peptide was added, and then the samples were incubated overnight. The eluates were analyzed by immunoblotting, Coomassie blue staining, and silver staining. The eluates were digested and subjected to NanoLC-MS/MS analysis. ProLuCID was used to identify the peptides (Xu, Park et al., 2015).

### Gene ontology analysis

ADCK4 interactors were analyzed using the Database for Annotation, Visualization and Integrated Discovery (DAVID) for functional annotation. The Functional Annotation Tool in the online version of DAVID (version 6.8) was run (http://david.abcc.ncifcrf.gov/) using the default parameters and gene ontology categories representing molecular function, cellular component, and biological process were separately analyzed for enrichment. *p* value < 0.05 was considered significant.

### Statistics

Statistical analyses were performed using Graph Pad Prism 7^®^ software. The results are presented as mean ± standard error or standard deviation for the indicated number of experiments. Statistical analysis of continuous data was performed with two-tailed Student’s *t* test or multiple comparison, as appropriate. Specific tests performed in the experiments are indicated in the figure legends. The results with *p* < 0.05 were considered statistically significant.

## Acknowledgments

We thank Dr. Jin Young Kim and Gina Yoon (Korea Basic Science Institute, Ochang, Division of Biomedical Omics Research) for the NanoLC-MS/MS analysis. We acknowledge the support of the UAB/UCSD O’Brien Core Center for Acute Kidney Injury Research for the LC-MS/MS analysis in this study. We also thank Maria Ericsson, Louis Trakimas, Elizabeth Benecchi, and Peg Coughlin from the Electron Microscope Core Facility, Harvard Medical School, for excellent TEM services and Evelyn Flynn for her outstanding technical assistance. We thank Yonsei Advanced Imaging Center for assistance with the Carl Zeiss microscope. This study was supported by the National Institutes of Health to FH (DK076683). F.H. is the William E. Harmon Professor. H.Y.G. was supported by the Chung-Am (TJ Park) Science Fellowship and the Research Program through the National Research Foundation of Korea (NRF) funded by the Korea Government (MSIT, 2018R1A5A2025079). EW was supported by the Leopoldina Fellowship Program, German National Academy of Sciences Leopoldina (LPDS 2015-07).

## Author contributions

E.W., S.Y., M.N., H.H., D.S., W.I.C., and M.G.L. carried out the animal experiments. E. W., S.Y., A. N., Y.W.C., M.N., H.H., F.B, D.S., W.I.C., J.K., J.H.R., M.G.L., and C.F.C. carried out the cell experiments. E.W., S.Y., F.H., and H.Y.G. conceived and directed the study. E.W. and S.Y. wrote the paper with help from F.H. and H.Y.G. The manuscript was critically reviewed by all the authors.

